# Quantification of the volume of swallowed air in the gut finds low volumes when asleep may reduce aerobic digestion and explain why short dinner to sleep times are associated with nocturnal reflux

**DOI:** 10.1101/2025.01.16.633483

**Authors:** Thomas Hurr

## Abstract

It has been previously reported that air swallowing and breathing exercises could reduce the severity of digestive reflux by supplying oxygen directly to the gut lumen and supporting aerobic digestion, however the normal volume of air swallowed over 24 hours has not been determined. To determine the volume of air swallowed over 24 hours, the number of swallows during eating, drinking and snacks (EDS), asleep, at other times awake (OTA) and the volume of air swallowed per bolus were sought from the literature. Four models were developed to determine the volume of air swallowed per bolus, finding volumes between 0 ml and an average maximum pharyngeal volume of 40 ml were possible, with an average and range of values ≈ 11(1.7-32) ml. From a literature search, the number of swallows over 24 hours determined using a microphone, was found to be the most complete set of data to calculate the volumes of air swallowed while EDS, asleep and OTA. There was on average during EDS ≈ 31 ml air swallowed per minute, when asleep ≈ 1 ml air swallowed per minute and at OTA ≈ 4.3 ml air swallowed per minute giving a total air swallow volume of ≈ 6,400(320-47,000) ml air over 24 hours. The volume of the gases contained in swallowed air were also calculated as nitrogen ≈ 5000 ml, oxygen ≈ 1000 ml and noting swallowed air is expired air from the lungs, carbon dioxide ≈ 320 ml over 24 hours. If improved aerobic digestion reduced the probability of digestive reflux and was related to the volume of air swallowed, then digestive reflux would be least likely to occur during EDS, with the highest air swallow rate, followed by OTA and most likely to occur when asleep, when the lowest volume of air is swallowed. The volume of air swallowed over 24 hours was equivalent to only one or two minutes of breathing at ≈ 6,000 ml per minute for an adult at rest. It is still not clear whether luminal oxygen supply from air swallowing, or luminal (and systemic) oxygen supply from breathing, is the major source of oxygen supply to the gut lumen for aerobic digestion, however if air swallowing is the major source of luminal oxygen supply, then air swallowing is likely an important factor for digestive health.

**Graphical Abstract:** 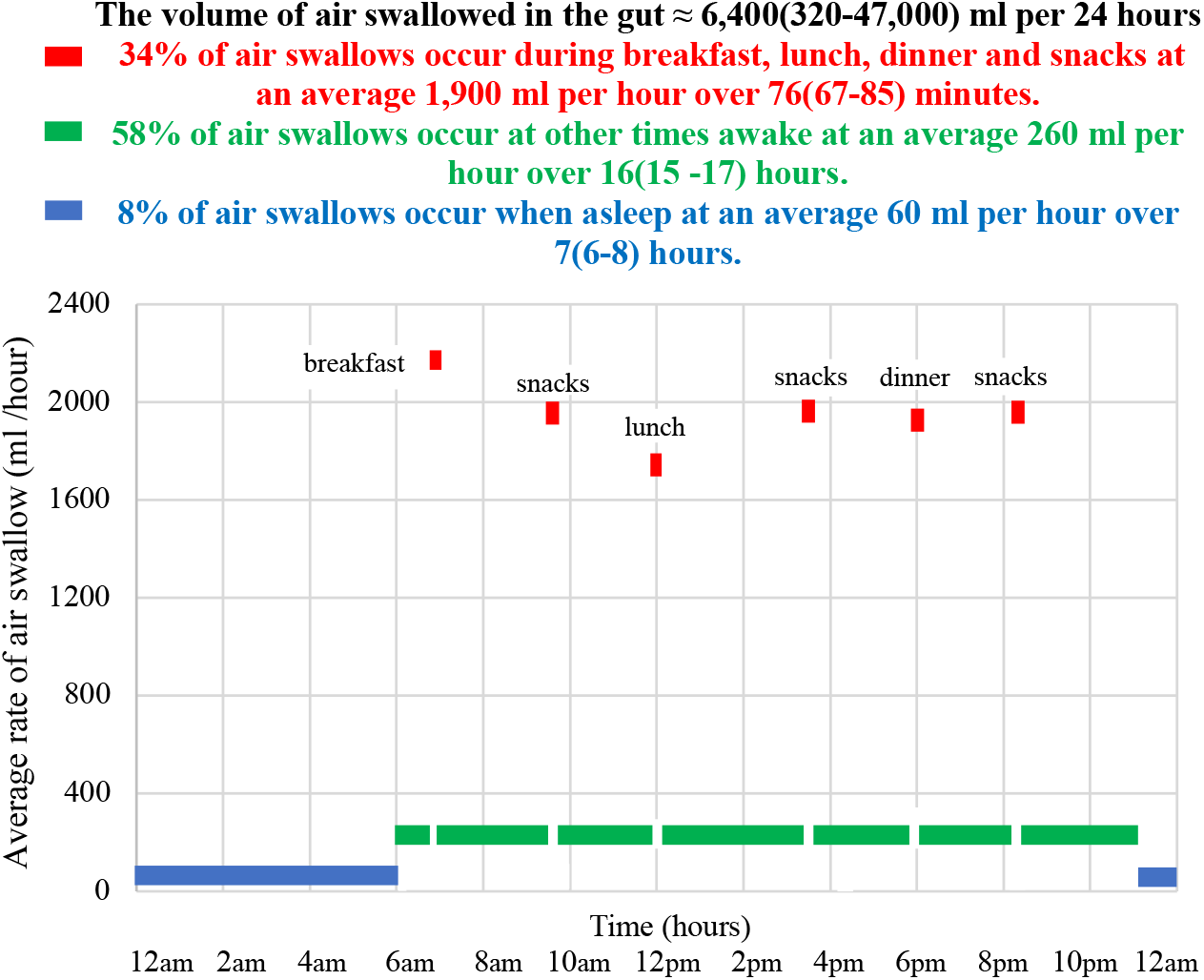

The low volumes of air swallowed during sleep may reduce aerobic digestion and explain why eating, drinking and snacks less than 3 hours before sleep have been associated with an increased probability of gastric /digestive reflux.

The volume of air swallowed per 24 hours is equivalent to only 1-2 minute of breathing at 6,000 ml per minute for an adult at rest.

Air swallowing link to reflux diseases [8], air swallow volume per bolus [9-14], rates of swallowing per 24 hours [16] breathing air volumes [35], increased reflux less than 3 hours dinner to sleep [38].

## 1. Introduction

Swallowing has been described as the movement of substances from the mouth to the stomach via the pharynx and esophagus [1]. Air swallowing can occur when eating, drinking or snacking (EDS), between meals and at other times awake (OTA) and asleep [2]. Excessive air swallowing or aerophagia has been associated with belching, bloating and dyspepsia [2-7]. Increased air swallowing has been reported for patients with dyspepsia and gastroesophageal reflux disease (GERD) refractory to proton pump inhibitor (PPI) and it has been suggested, may be a contributing factor [5-6]. Normal air swallowing has been considered essential in promoting beneficial aerobic digestion by direct chemical oxidation of food in the gut lumen and supporting the microbiome, reducing the probability of developing digestive reflux diseases [8].

This report quantifies the normal volume of swallowed air while EDS, OTA and asleep, as a starting point towards understanding the role of air in the digestive system.

Firstly, this study develops four models to quantify the possible range of values for the volume of air swallowed /bolus. Secondly, five different methods, that have been used to determine the number of swallows while EDS, OTA and asleep are reviewed, with calculated values from food volumes and bolus sizes, to determine the volume of swallowed air /24 hours. Thirdly, the movement of swallowed air in the gut is discussed and finally the volume of gases, that make up the composition of swallowed air, are calculated. The relevance of air swallowing to digestive reflux diseases is also considered.

## 2. Results

### 2.1 Four models for the volume of air swallowed per bolus

A bolus can be considered a small round mass of substance especially chewed food, that is swallowed. In this report, a bolus can consist of either a mass of solid or liquid food, saliva swallowed without food and air swallowed without food or saliva, as they all have mass and can be swallowed.

The volume of air swallowed per bolus (V_AS/B_) for 8 subjects using computer tomography (CT) scans of swallows with barium sulphate suspensions reported that 1 ml, 5 ml, 10 ml and 20 ml swallows resulted in pharyngeal chamber volumes (V_PHA_) of V_PHA_ ≈16, 20, 26, and 31 ml respectively [9,10]. Air could escape during the initial part of a swallow from the pharynx regardless of the bolus volume, resulting in ≈ 15 ml of air was swallowed for bolus volume 1-20 ml [9,10]. When the pharyngeal chamber was not occupied by a test bolus and presumably contained saliva, with the normal salivary rate between 0.5-1.5 ml /minute, the small bolus volume swallows were associated with substantial aerophagia [10]. The bolus after entering the pharynx was found to be randomly distributed around the perimeter, rather than contained as a single coherent bolus and mixed with a substantial quantity of air [9].

The volume of the pharynx (V_PHA_) from CT scans, had a median at rest (AR) range value of V_PHA_(AR) ≈20.1(16.8-26.0) ml. During the bolus swallowing process, the AR volume changes with tongue loading to V_PHA_ ≈20.1(11.6-38.6) ml, either decreasing due to escaping air, or increasing with bolus volume, prior to decreasing from pharyngeal contraction on swallowing to V_PHA_ ≈ 0-1ml, before taking in air and returning to the AR volume [11]. The volume of the whole pharynx did not exceed a maximum value of V_PHA_(Max)≈ 38.6 ml [11].

Another study reported the volume of air swallowed by 7 subjects with swallows of bolus volume (V_B_) as V_B_ ≈ 10 ml of barium contrast solutions, using CT scanning, that the resulting air swallow volume V_AS/B_ ≈ 17.7(8-32) ml [12]. For V_AS/B_ ≈ 32 ml and V_B_ ≈ 10 ml, the value for the maximum pharyngeal volume is presumably V_PHA_(Max) ≈ 32 +10≈ 42 ml [12]. The maximum value from the 2 studies can be averaged to give V_PHA_(Max) ≈ (38.6 + 42)/2 ≈ 40 ml [11,12].

The release of air from the pharynx as part of the swallowing process, has been reported in another study where, regardless of bolus volume, with V_PHA_ ≈ 20 ml before a swallow, ≈16 ml of air escapes, resulting in a low air swallow volume /bolus of V_AS/B_ ≈ 2.9(1.7-4.0) ml [13].

A study of the effects of bolus consistency on pharyngeal volume during swallowing found for thin barium swallow solutions ≈ 8 ml was swallowed and ≈ 12 ml of air was swallowed. For thick barium swallow solutions ≈ 7.1 ml was swallowed and ≈ 5.3 ml air was swallowed suggesting the volume of an air swallow can be influenced by the solution density [14]. These results gave an average for V_AS/B_ ≈ (12+5.3)/2 ≈ 8.6 ml [14].

Four models to describe the possible volume of air swallowed /bolus are presented.

Model 1. If the maximum volume of the pharynx is ≈ 40 ml and can contain 40 ml of air, then on swallowing a bolus, the volume of air equivalent to the volume of the bolus must escape to prevent exceeding the maximum value resulting in a reduced air swallow volume:

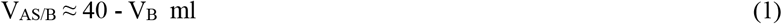

and could result in very high air swallow volumes (≈ 40 ml) at low bolus volumes and gives the maximum values for air swallows /bolus.

Model 2. The at rest pharyngeal volume of V_PHA_(AR) ≈ 20 ml, but with the pharynx able to expand to a maximum volume of 40 ml, when a bolus is swallowed, no air needs to escape, if the bolus volume is < 20 ml, resulting in an air swallow volume of ≈ 20 ml, independent of bolus volume < 20 ml. When the bolus volume is >20 ml, air must escape as the maximum pharyngeal volume cannot exceed 40 ml, resulting in an air swallow volume <20 ml when V_B_ > 20 ml:

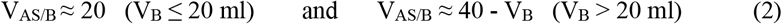

Model 3. If any quantity of air can escape from the pharynx during a swallow except for a residual amount, then the air swallow volume becomes independent of bolus volume and a smaller air swallow volume can result, as has been found, with average and range of values reported [13]:

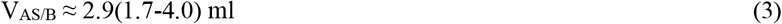

Model 4. Liquid only swallows have been reported for presumably small bolus volumes of saliva with all the air escaping during the swallow [2,3,5,7]:

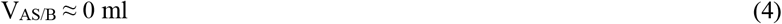

Which of the 4 models could be most applicable may depend on the bolus consistency, volume or whether in the states of EDS, OTA or asleep, Fig. 1.

**Figure 1.**
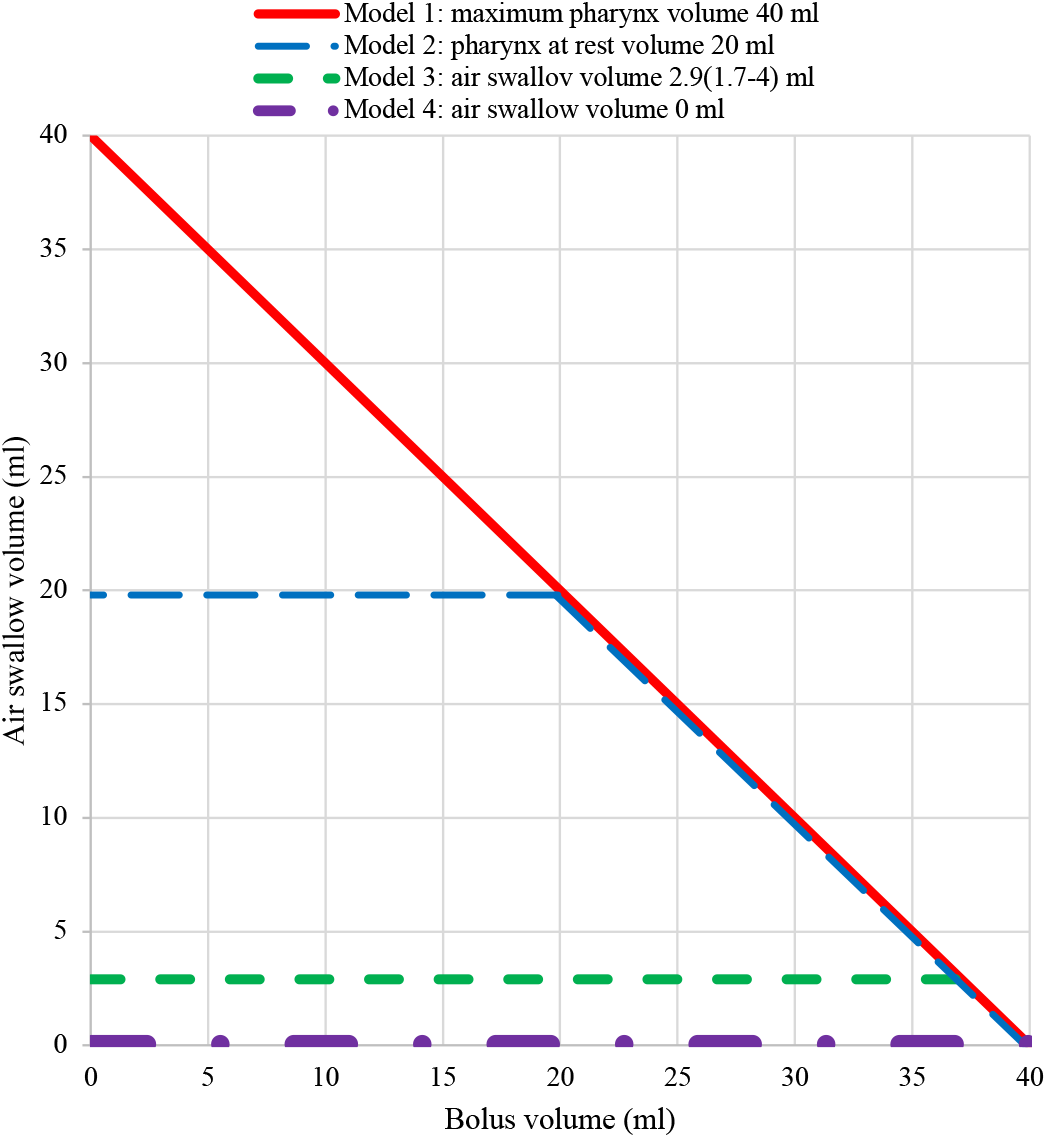
Four models for pharyngeal air swallowing (eqs. 1-4). For the first model, the pharynx has a maximum volume of air ≈ 40 ml when the bolus volume V_B_ ≈ 0 ml, but on swallowing a bolus, air equivalent to the bolus volume must be released resulting in a lower air swallow volume (eq.1) and when V_B_ ≈ 40ml, the air swallow volume is ≈ 0 ml. For the second model the pharynx has an at rest volume of ≈ 20 ml and when the bolus volume V_B_ < 20 ml, an air swallow volume of up to 20 ml can occur but when V_B_ > 20 ml, air must be released resulting in air swallow volumes < 20 ml (eq.2). For the third model where most air can escape the pharynx V_AS_ ≈ 2.9(1.7-4.0) ml and essentially independent of V_B_, as has been reported [13]. For the fourth model, all air escapes before a swallow, reported as a liquid only swallow from impedance measurements within the oesophagus, then V_AS_ ≈ 0 ml [2,3,5,7].

From the 4 models, V_AS/B_ ≈ 11(0-40) ml with V_AS/B_ ≈ 0 for liquid only swallows. It has been reported that most swallows performed during meals were found to contain gas (95±1%) as well as outside of meals (99±1%) suggesting V_AS/B_ ≈ 0 ml is uncommon and so outside the normal range of values [4]. Similarly, V_AS/B_ ≈ 40 ml is too large and also considered outside the normal range. The air swallowed per bolus from radiographic studies using barium swallows, reported V_AS/B_ ≈ 15 ml, 17.7(8-32) ml, 2.9(1.7-4.0) ml, 8.6(5.3-12) ml, and were used to generate an average and range of values V_AS/B_ ≈ 11(1.7-32) ml, less than a third of the maximum pharyngeal volume, V_PHA_(Max) ≈ 40 ml with range, the smallest and largest air swallow volumes given [9-14].

### 2.2 Duration of time in the states of EDS, OTA and sleep

The number of swallows /minute (N_S_/min) can be calculated from the number of swallows in the state (N_S_(state)), in minutes, over the time in the state, t(state) in minutes, making the time in each state an important factor in determining swallow rates:

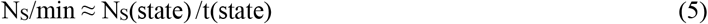

The duration of mealtimes /24 hour for 10 patients with excessive belching and 11 people in the control group were both found to be almost the same times, with all mealtimes added together giving an average of ≈ 76±6 minutes /24 hours [4]. For 44 patients with GERD included 18 patients responsive to proton pump inhibitors (PPIs) and 26 patients unresponsive to PPIs, were similarly almost the same times with the average of all mealtimes added together ≈ 83±9 minutes/24 h [15]. A study of swallowing for 20 healthy adolescents and young adults found average mealtimes ≈ 70 minutes [16]. From the 3 different studies, the average time in the state of EDS (presumably actively swallowing food) ≈ (76+83+70)/3 ≈ 76±9 minutes /24 hours or ≈ 1 hour 16±9 minutes (1.27 hours) /24 hours [4,15,16]. The time EDS does not occur when asleep and therefore over less than 24 hours but is often written as EDS /24 hours.

A national survey with 37,832 people who completed interviews based on recall, found people aged 15 years or older had an eating or drinking time ≈ 67 minutes as the primary or associated primary activity and while doing another activity ≈ 78 minutes adding to ≈ 2.5 hours /24 hours eating or drinking, with eleven percent of the population spending at least 4.5 hours on average engaged in eating and drinking activities [17]. These times are significantly greater than the ≈ 76 minutes discussed above, presumably as time eating and drinking at meals includes time doing other activities, including social activities. The use of the term prandial presumably includes both the times actively eating, drinking and snacks and time doing other things including social activity during the eating drinking and snack time. In this report the three different states were divided into EDS, asleep and OTA. Use of the terms eating and drinking, sleeping and with other activities to describe the 3 different states has been used previously, possibly to avoid any confusion with the use of the term prandial [16].

Polysomnography data from 206 healthy subjects of aged 19-73, found the average time asleep for the first night was 6 hours 32 minutes (5 hours 42 minutes to 7 hours 23 minutes) which changed after the first night giving for the second night 7 hours (6 hours 23 minute to 7 hours 34 minutes) [18]. Another study with 2838 adult participants found the average self-reported sleeping time was ≈ 7 hours (no range of times given) [19]. From these 2 studies, the average and range of time asleep in hours is estimated to be ≈ 7(6-8) hours /24 hour [18,19]. A calculated time in hours for OTA ≈ 24-1.27-7(6-8) ≈ 16(15-17) hours giving the average times for the 3 states /24 hours: EDS ≈ 1.27 hours (76±9 minutes), asleep ≈ 7(6-8) hours and OTA ≈ 16(15-17) hours /24 hours.

### 2.3 Method 1: the number of swallows determined by microphone

An article from the literature using measurements with a microphone, placed on the neck of 20 healthy adolescent and young adults, who were allowed to remain in their homes over 24 hours, determined the number of swallows (N_S_) in each of the states of EDS (70 minutes), asleep 8.5 hours (510 minutes) or OTA 14.3 hours (860 minutes) /24 hours (eq. 5), Table 1, 2 [16]. Sleeping times were 1.5 hours longer than the average values above, possibly due to the younger age of the study group [16]. The number of swallows in each state (N_S_(state)) can been multiplied by the volume of air swallowed /bolus (V_AS/B_) to determine the volume of air swallowed (V_AS_) in each state (V_AS_(state)) :

**Table 1.** The number of swallows in each state as eating drinking snacks (EDS) ≈ 200(80-510) /70 minutes for EDS /24 hours, asleep ≈ 45(19-90) /510 minutes for asleep /24 hours and at other times awake (OTA) ≈ 340(86-589) /860 minutes for OTA /24 hours with the range of air swallow volumes /bolus of ≈11(1.7-32) ml to give a calculated average and range of values for the volumes of air swallowed in each state /24 hour, /hour or /minute and /hour assuming air is swallowed with each swallow. From the total number of swallows, the % swallows in each state EDS ≈200/585≈ 34%, asleep 45/585 ≈ 8% and OTA ≈ 340/585 ≈ 58% can be calculated. Calculation methods in Appendix 1.

| State [16] | Number of swallows 585(185-1459) /24 hours [16] | Number of swallows /minute (eq.5) [16] | Volumes of air / swallow ml /24 hours (eq.6) | Volume of air swallow ml /hour (eq.7 x 60) | Volume of air swallow ml /minute (eq.7) |
| --- | --- | --- | --- | --- | --- |
| Eating, drinking, snacks 70 minutes | 200(80-510) | 2.9(1.1-7.3) | 2,200(140-16,000) | 1,900(120-14,000) | 31(1.9-230) |
| Asleep 8.5 hours | 45(19-90) | 0.087(0.037-0.18) | 491(32-2,900) | 60(3.8-340) | 0.96(0.063-5.7) |
| Other times awake 14.3 hours | 340(86-859) | 0.39(0.10-1.0) | 3,700(150-27,500) | 260(10-1,900) | 4.3(0.17-32) |

**Table 2.** Number of swallows /24 hours and /minute with the volume of air swallowed /24 hours, /hour and /minutes in all states summed together. Calculation methods in Appendix 1.

| Number of swallows /24 hours in all states added together [16] | Number of swallows /minute in all states [16] | Volumes of air /swallow (ml) /24 hours, in all states (eq. 6) | Volume of air swallow ml /hour in all states ranges (eq. 6, /24) | Volume of air swallow ml /minute in all states (eq. 6, /24x60) |
| --- | --- | --- | --- | --- |
| 585(185-1459) | 0.41(0.13-1.0) | 6400(320-47,000) | 270(13-2,000) | 4.5(0.22-32) |

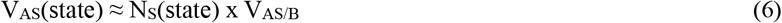

To calculated volume of air swallowed /minute in each state, eq. 6 is divided by the t(state) in minutes:

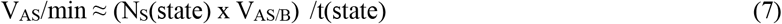

To determine the average volume of air swallowed /minute (V_AS_/min), the average number of swallows in each state was multiplied by the average volume of air swallow /bolus (≈ 11 ml) and divided by the time in each state, t(state). For the lowest volume of air swallowed, the lowest number of swallows in each state and the lowest air swallow volume (≈ 1.7 ml) and for the highest value, the highest number of swallows in each state and the highest air swallow volume (≈32 ml), divided by the time in the state, to give the average and range of air swallow volumes /24 hours, Table 1, 2. To convert the air swallow from ml /minute to ml /hour, values from eq. 7 are multiplied by 60, Table 1.

The swallow rates per minute (Table 1) can be converted to the number of minutes between swallows giving 1/2.9 ≈ 0.34 minutes /swallow while EDS, 1/0.087≈ 11.5 minutes /swallow asleep and ≈1/0.39 ≈ 2.6 minutes /swallow when at OTA.

The article from the literature also gave the number of swallows during breakfast, lunch, dinner and snacks which on multiplying by V_AS/B_ ≈ 11ml can give the V_AS_ / min or /hour (eqs. 6,7) for:

breakfast; 37.6 swallows over 10.9 minutes giving ≈ 3.4 swallows /minute or 207 swallows /hour or ≈ 2280 ml air /hour,

lunch; 37.3 swallows over 14.2 minutes giving ≈ 2.6 swallows /minute or 158 swallows /hour or ≈ 1730 ml air /hour,

dinner; 64.2 swallows over 22.6 minutes giving ≈ 2.8 swallows /minute or 170 swallows /hour or ≈ 1870 ml air /hour,

snacks; 65.2 swallows over 22.5 minutes giving ≈ 2.9 swallows /minute or 174 swallows /hour or ≈ 1912 ml air /hour [16].

The results show a faster rate of swallows /minute at breakfast compared to a higher number of swallows for dinner and snacks but over longer times, resulting in slower swallow rates (air swallow rates /hour shown in the graphical abstract) [16].

Different times for eating breakfast 17.1±7.9 minutes, lunch 27.5±10 minutes, dinner 24.4±10.9 minutes and snacks 11.8±7.4 minutes have been determined for children aged 5 years old, with eating time ≈ 80.8±27.3 minutes [20].

Use of a microphone was thought the least invasive technique that most simulated normal human conditions /24 hours and selected to calculate the air swallow volume /24 hour, despite differences in the average times EDS, asleep and OTA as discussed above, Table 1, 2 [16].

### 2.4 Method 2: the number of swallows during sleep determined by polysomnography

A study using polysomnography and surface electromyography of swallowing during sleep with 10 adults aged 71-81 years found the median number of swallows during sleep were ≈ 0.068(0.005-0.093) /minute, with swallow free periods up to 70.5 minutes [21]. The values are lower than the values of ≈ 0.087(0.036-0.18) /minutes reported for the 20 young adult subjects using a microphone, about 60 years ago, with swallow free periods of only 20 minutes or more, during sleep, Table 1 [16]. Swallow rates during sleep clearly decline with age [16,21].

### 2.5 Method 3: the number of swallows determined by impedance measurements

Impedance measurements can also determine the number of swallow /minute using a catheter placed within the oesophagus and distinguish the type of swallow, as liquid only swallows, air swallows and liquid with air swallows over time, reported in the states of prandial, supine, asleep or pre /postprandial [2-5]. Most swallows performed during meals were found to contain gas (95±1%) as well as outside of meals (99±1%), with swallow rates decreasing when supine, Table 3 [4]. Values for the number of air swallows /minute for both healthy controls and patients with belching, belching /aerophagia, dyspepsia, GERD and GERD/dyspepsia who have undergone treatment with proton pump inhibitors (PPIs) have been reported. Swallows from only the healthy control groups from these reports are shown in Table 3. The average values of total swallows pre /postprandial swallows ≈ 1.09(0.20-1.73) /minute are higher than the value for prandial swallows 0.9(0.73-1.1) /minute with both much higher than when supine and awake ≈ 0.05±0 /minute or when asleep ≈ 0.042±0.002 /minute, Table 3.

**Table 3.**
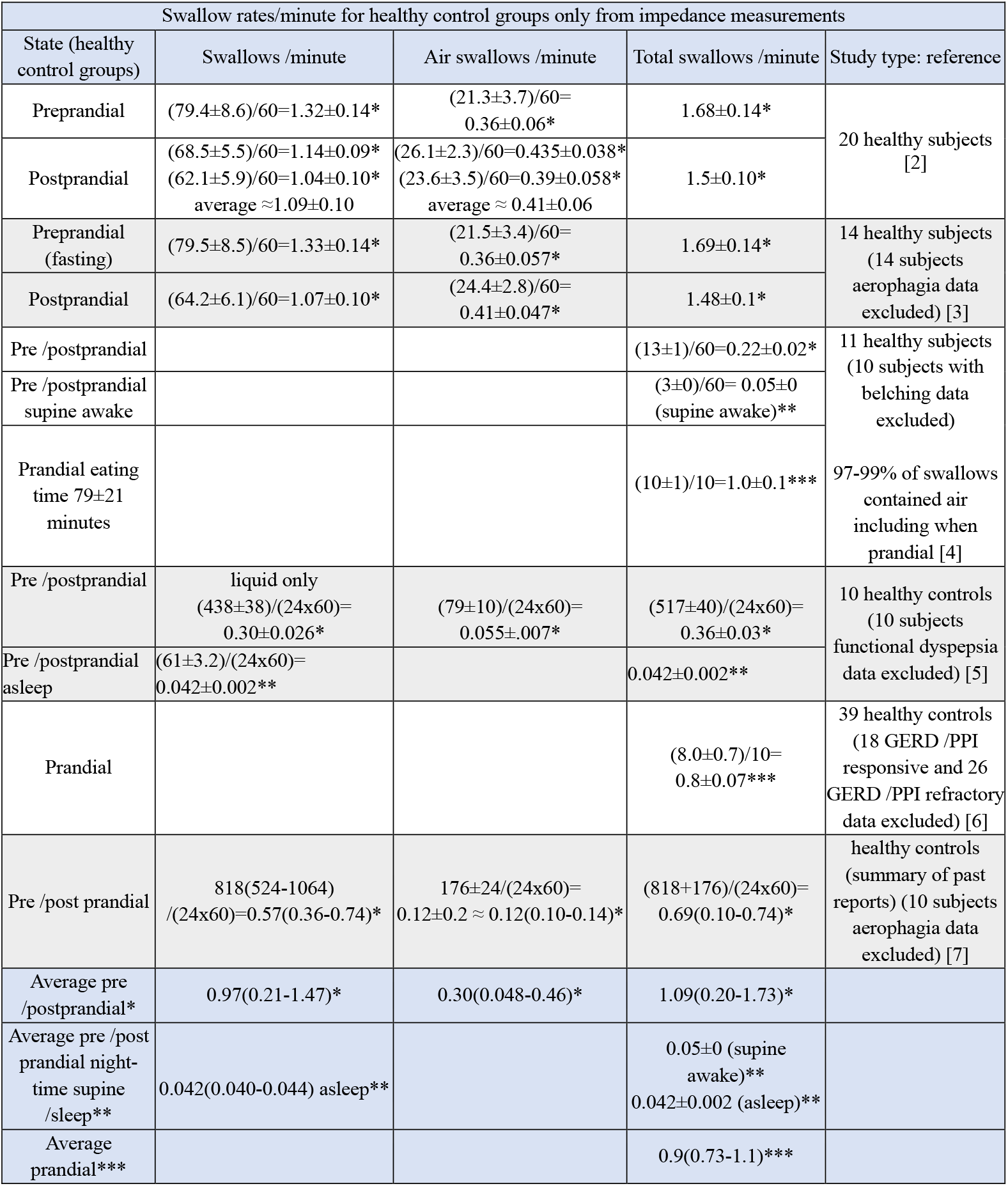
Swallow and air swallow rates (ml /minute) determined from impedance measurements for healthy control subjects. To determine an average value for air swallows /minute for the 3 states, values for the pre /postprandial*, pre /postprandial night-time supine or asleep** and prandial*** as indicated, were averaged in each column. The swallow values were reported as swallows /24h, /hour or /10 minutes and converted swallows/minute. The range of values was simply taken as the smallest and largest values possible in each set of numbers used to form the average values. Gaps in the table where no data was available.

### 2.6 Method 4: the number of swallows calculated from food and bolus volumes

To quantify the volume of air swallowed during a meal, the number of bolus (N_B_) swallowed and the volume of air swallowed /bolus (V_AS/B_) are required. To determine the number of boluses swallowed, the volume EDS as (V_EDS_) and the bolus volume (V_B_) are required:

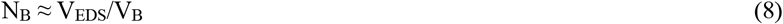

which when multiplied by the volume of air swallowed /bolus (V_AS/B_) gives the total volume of air swallowed (V_AS_):

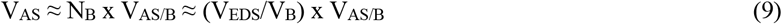

The volume of food and beverages consumed each day (the same value over 24 hours) from a national survey (24,000 people) found V_EDS_ ≈ 3,100g (range 3,000-3,700 g) /person /24 hours [22]. If it is assumed the density of both the solid food (usually containing water) and liquid foods, are comparable to that of water, then 3,100(3,000-3,700) g /24 hours ≈ 3,100(3,00-3,700) ml /24 hours allowing values to be measured in volume.

In another study, the volume of liquid (water, beverages and food moisture) from 22,716 children and adults estimated the volume to be ≈ 2,700 ml, consumed per person /24 hours [23]. From the average volumes of food consumed V_EDS_ ≈ 3,100(3,000-3,700) ml /24 hours and liquid consumed ≈2,700 ml, then the volume of solid food can be calculated as ≈ 3,100 - 2,700 ml or ≈ 400 ml /24 hours.

To quantify average bolus volumes, a study with 95 volunteers using barium contrast swallows and CT scans found the bolus volume of self-selected swallows, defined as a swallow volume based on what is most comfortable to swallow in full, had an average bolus volume V_B_ ≈ 16.66 (8.96 - 24.36) ml [24]. Another study with 38 healthy adults using videofluoroscopy found that sips of thin liquid barium swallows had a V_B_ ≈12.13±6.68 ml decreasing for thick liquids to V_B_ ≈ 5.15±2.59 ml [25].

From the volume of food V_EDS_ ≈ 3,100(3,000-3,700) ml /24 hours and assuming a self-selected bolus volume V_B_ ≈ 16.66(8.96-24.36) ml, from eq. 8 the average value for N_B_ ≈ 3100/16.66 ≈ 186 with the lowest bolus number occurring when N_B_ ≈ 3000 /24.36 ≈123 and the highest bolus number when N_B_ ≈ 3700 /8.96 ≈413, give an average and range for N_B_ ≈186(123-413) /24 hours which occur when EDS.

For V_EDS_ ≈ 3,100(3,000-3,700) ml /24 hours, self-selected bolus volume, V_B_ ≈ 16.66(8.96-24.36) ml and the volume of air swallowed per bolus V_AS/B_ ≈ 11(1.7-32) ml, the average volume of air swallowed (V_AS_/24h) can be calculated from eq. 9 as:

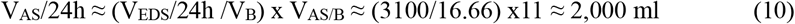

To determine the lower value for V_AS_/24 hours, the least volume of food consumed V_EDS_ ≈ 3000 ml larger self-selected bolus volume V_B_ ≈ 24.36 ml and least volume for air swallowing /boluses V_AS/B_ ≈ 1.7 ml, gives a value for V_AS_/24 hours ≈ 210 ml. To determine the higher value for V_AS_/24 hours, the largest volume of food consumed V_EDS_ ≈ 3,700 ml, the smallest self-selected bolus volume V_B_ ≈ 8.96 ml and the largest volume for air swallowing/bolus V_AS/B_ ≈ 32 ml gives a large value of V_AS_/24 hours ≈ 13,200 ml. The resultant average and range for the volume of air swallowed, V_AS_/24 hours ≈ 2,000(210-13,200) ml while EDS over 24 hours, and is comparable to the value of 2,200(140-16,000) calculated from the number of swallows while EDS, Table 1.

The number of swallows /min over the time EDS in minutes (tEDS) can be calculated from eq. 8 as:

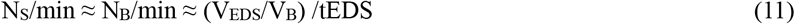

where tEDS ≈ 76(70-83) minutes. For meal volumes of V_EDS_ ≈ 3,100(3,000-3,700) ml, self-selected food bolus volume V_B_ ≈ 16.66(8.96-24.36) ml and tEDS ≈ 76(70-83) minutes, the average number of swallows /minute (N_S_ /min):

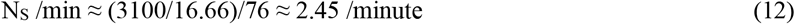

To generate the lowest value for N_S_ /min, the least volume of food consumed V_EDS_ ≈ 3,000 ml, the largest value for the bolus volume V_B_ ≈ 24.36 ml and the longest eating time of 83 minutes gives N_S_ /min ≈ (3000/24.36)/83 ≈ 1.48 /minute. To generate the largest value for N_S_ /min, the largest volume of food consumed V_EDS_ ≈ 3,700 ml, the lowest value for bolus volume V_B_ ≈ 8.96 and the least value for eating time of 70 minutes gives N_S_ /min ≈ (3700/8.96)/70 ≈ 5.9 /minute. The resultant average and range of N_S_ /min ≈ 2.5(1.5-5.9) minute /≈ 76(70-83) EDS /24 hours for V_EDS_ ≈ 3,100(3,000-3,700) ml.

### 2.7 Method 5: the number of saliva swallows and swallow volume from counting sensors and weight

Saliva swallow rates can be determined from several methods including direct observation, self-counting, sound detection using microphones and sensors [26]. Saliva swallows have been described as either voluntary occurring especially at mealtimes or spontaneous swallows usually unconscious and without purposeful intervention when asleep and OTA [26,27]. Saliva swallows have also been described as stimulated swallows while eating or chewing gum and undertaking a saliva sampling test, and unstimulated swallows at times other than eating or chewing gum or undertaking saliva sampling tests [28,29]. Saliva has a major role in providing bicarbonate ions for buffering acid in the stomach and esophagus and the physical clearance of the esophagus [27-29]. Note the large decline in saliva swallow rates with age decreasing from ≈ 0.98 /minute to ≈ 0.21 /minute, Table 4 [26]. Saliva swallow rates were not found reported in the state of EDS but saliva production rates while prandial for children, but not for adults, were found, Table 5.

**Table 4.** Saliva swallow rates for the healthy control groups are ≈0.98(0.47-1.7) /minute <60 years old, decreasing with age to ≈0.21(0.13-0.47) when >60 years old [26]. *a swallow interval was given as 60.8±39 sec and recalculated as swallows /minute ≈ 60/60.8(60/99.8-60/21.8) ≈0.99(0.6-2.75) /minute [30]. ** Values were highly variable and differed greatly among individuals, calculated value from the number of swallows as 119.1(12-457) /time asleep as 420(203-551) minutes giving a calculated average of 119.1/420≈ 0.28 /minute, a minimum of 12/551 ≈ 0.022 /minute and a maximum range 457/203 ≈ 2.25 /minute or 0.27(0.022-2.25) swallows /minute over 203-551 minutes [27].

| Healthy controls: State | Number of saliva swallows /minute healthy controls | Study type: reference |
| --- | --- | --- |
| Awake (voluntary unstimulated swallows) | 0.98(0.47-1.7) < 60 years old<br>0.21(0.13-0.47) > 60 years old<br>average 0.60(0.13-1.7) | review, 19 articles with healthy controls (patients with dysphasia, Parkinson's disease excluded) [26] |
| Awake between 3-6 pm for 30 minutes (voluntary unstimulated swallows) | 0.99(0.60-2.75)* | 128 healthy volunteers, [30] |
| Asleep average 7 hours 17 minutes (spontaneous swallows) | 0.28(0.022-2.25)** | 18 healthy controls (21 patients with Parkinson's disease excluded) [27] |

**Table 5.** Saliva volume /minute awake unstimulated, awake stimulated by saliva testing for patients using polypharmacy, awake stimulated by chewing gum for GERD patients who had used PPIs and for children aged 5 when chewing food including when awake and not eating. Greatest saliva production occurs when eating food followed by awake stimulated (but not eating), awake and unstimulated.

| State | Saliva production volume ml /minute | Study type: reference |
| --- | --- | --- |
| Awake (voluntary unstimulated) | 0.56± 0.41 | 128 healthy volunteers, [30] |
| Awake between 7.30 am and 5.30 pm (voluntary stimulated by saliva collected) | 0.967± 0.433 | 6753 patients using polypharmacy, [29] |
| Awake stimulated by sugarless chewing gum (voluntary stimulated) | 1.3(0.7-2.0) | 32 GERD /PPI patients, [28] |
| Chewing foods for $\approx 80 \pm 27$ minute, (voluntary stimulated) | 3.6(2.8-4.4) | 30 children aged 5, total saliva volume 288 ml (chewing food) + 208 ml (awake not eating) assuming almost no saliva produced during sleep $\approx 500$ ml /24 hour [20] |
| Awake not eating (voluntary unstimulated, $\approx 13.5$ hours) | 0.26(0.1-0.42) | |

Saliva swallow volumes have been determined by expelling saliva into a container, placing gauze in the mouth, or chewing food and spitting out before swallowing and measuring the change in weight over time, Table 5 [20,28-30]. For 128 healthy volunteers the number of awake voluntary unstimulated saliva swallows ≈ 1 /minute with a saliva production volume ≈ 0.56 ml suggesting a saliva bolus has a swallow volume of ≈ 0.56 ml /minute at OTA Table 4, 5 [30].

An approximation of the volume of saliva produced by adults while awake and unstimulated for an average 16 hours presumably as OTA, can be calculated from the average unstimulated saliva production of 0.56 ml as ≈0.56(16×60) ≈ 538 ml saliva. Assuming the child and adult saliva production while eating was approximately equal at ≈ 288 ml (values for adults not found available) and with very little saliva produced while asleep (values while asleep not found available), values for OTA ≈ 538 +288 ≈ 830 ml /24 hours for healthy adults, with larger values predicted for the polypharmacy and GERD patient groups and a decline in values with age, Table 5 [20,26,30]. It has been reported for 5 years old children stimulated and unstimulated saliva production was ≈ 500ml /24, Table 5 [20]. More data is required to improve the accuracy for the calculation of saliva production while EDS, asleep and OTA to determine the total volume /24 hours.

### 2.8 A comparison of the 5 methods to determine the number of swallows per minute

The number of swallows /minute determined from the five different methods, including in the states of OTA, asleep and EDS, from Tables 1-4 for healthy subjects are summarized in Table 6.

**Table 6.** The numbers of swallows including liquid and air swallows /minute in each state determined by 5 different methods for healthy control groups, Tables 1-4. The states were reported as prandial, pre /post prandial for the impedance and saliva measurements. * Values were highly variable and differed greatly among individuals [27]. Gaps in the Table where no data available.

| State<br>(healthy control groups averaged values where possible) | Method 1:<br>number of swallows /minute<br>microphone method [16] | Method 2: number of swallows /minute from polysomnography and electromyography 71-80 years old [21] | Method 3: number of swallows /minute from impedance measurements [2-7] |  |  | Method 4:<br>number of swallows /minute calculated from food /food bolus volumes [22-24, this work] | Method 5:<br>number of saliva swallows /minute from counting /sensors [26,27] |
| --- | --- | --- | --- | --- | --- | --- | --- |
|  |  |  | liquid | air only | liquid + air added together |  |  |
| OTA or pre /postprandial | 0.39(0.10-1.0) |  | 0.97(0.21-1.47) | 0.30(0.048-0.46) | 1.09(0.20-1.73) |  | 0.98(0.47-1.7) < 60 year old;<br>0.21(0.13-0.47) > 60 year old;<br>average 0.60(0.13-1.7) |
| Asleep | 0.087(0.037-0.18) | 0.068(0.005-0.093) |  |  | 0.042±0.002 |  | 0.28(0.022-2.25)* |
| Supine awake |  |  |  |  | 0.05± 0 |  |  |
| EDS or prandial | 2.9(1.1-7.3) |  |  |  | 0.9(0.73-1.1) | 2.5(1.5-5.9) |  |

Firstly, for OTA from the microphone method, the number of swallows /minute ≈ 0.39(0.10-1.0) /minute, was significantly lower than the average value from impedance measurements of ≈ 1.09(0.20-1.73) /minute and about two thirds the average value from saliva swallows ≈ 0.60(0.13-1.7) /minute, Table 6. For saliva swallows when < 60 years old, the swallow rate ≈ 0.98(0.47-1.7) /minute is similar to the average value measured from impedance of ≈ 1.09(0.02-1.73) /minute, reducing significantly by ≈ 4 times with age to ≈ 0.21(0.13-0.47) /minute > 60 years old, Table 6 [26]. The reduced saliva swallowing with age (> 60 years old) also suggests a decrease in air swallowing with age at OTA, possibly indicating an increased risk of digestive disease with age, Table 6.

Secondly, the number of swallows /minute while asleep for the microphone method, polysomnography and electromyography and while asleep or supine from impedance measurements were all relatively low and between ≈ 0.042-0.087 /minute, less than a third the value of ≈ 0.27(0.06-2.75) /minute for saliva swallows, Table 6. Values for saliva swallows, when asleep of ≈ 0.27(0.06-2.75) /minute were highly variable and differed greatly among individuals, requiring further investigation to determine what trends may exist for healthy control groups [27].

Thirdly, the number of air swallows /minute while EDS ≈ 2.9(1.1-7.3) /minute is approximately three times the value for the number of swallows while prandial (EDS) from impedance measurements ≈ 0.9±0.2 /minute, Table 6. The number of prandial swallows ≈ 1 swallow /minute over ≈ 80 minutes results in ≈ 80 swallows over 24/ hours and appears to be a low value. To determine the bolus volume using the value for N_AS_/min ≈ 0.9 /minute, (Table 6) the average value of EDS /24 hours of V_EDS_ ≈ 3100 ml and the average time prandial or EDS ≈ 76 minutes from rearranging eq. 11:

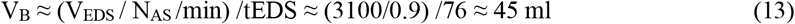

showing the bolus volume is too large and greater than V_PHA_(Max) ≈ 40 ml. To determine the bolus volume using the value for N_AS_/min ≈ 2.9 /minute, (Table 6) the average value of EDS /24 hours of V_EDS_ ≈ 3100 ml and the average time prandial or EDS ≈ 76 minutes from rearranging eq. 11:

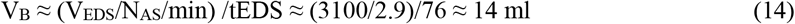

which is within the range of the self-selected bolus volumes, V_B_ ≈16.66 (8.96-24.36) ml [24]. The average swallow rates from impedance measurements when prandial ≈ 0.9±0.2 /minute are also essentially the same as the average pre /postprandial swallow rate ≈ 1.09(0.20-1.73) /minute, Table 6. The low number of swallows /minute while prandial may be due to a lower volume of food consumed during the measurements, or increased bolus volumes or tEDS, eq. 11.

For children 5 years old, stimulated saliva production when EDS (3.6 ml /minute) compared to unstimulated production at OTA (0.26 ml /minute) indicates 3.6/0.26 ≈ 14x more saliva produced when EDS, Table 5 [20].

The number of swallows /min while EDS for the microphone method ≈ 2.9(1.1-7.3) /minute were similar to the calculated value based on food and bolus volumes ≈2.5(1.48-5.9) /minute, Table 6.

### 2.9 The movement of swallowed air in the gut

A magnetic resonance imaging (MRI) study of the stomach found baseline gas volume of ≈27±14 ml increased on ingesting 500 ml of soup to an initial gas volume ≈109±55 ml, with little gas volume change after 60 minutes as ≈102±63 ml [31]. From the self-selected bolus volumes V_B_ ≈ 16.66(8.96-24.36) ml, using V_B_ ≈ 8.96 ml and a volume of swallowed air /bolus of V_AS/B_ ≈ 1.7 ml, from eq. 9:

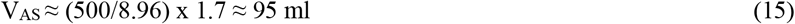

with the number of boluses swallowed N_B_ ≈ 500/8.96 ≈ 56 (eq. 8), consistent with the MRI result. If average self-selected bolus and air swallow volumes were used:

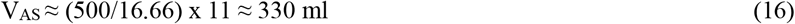

which is ≈ 220 ml larger than the value measured by MRI (102-109 ml) and could suggest ≈1/3 of swallowed air is retained during digestion and 2/3 expelled [24,31]. It is likely the volume of swallowed air is not represented by the volume of gas retained in the stomach over time, with excess gas eructated, oxygen used for aerobic digestion, gas absorption into tissue and the blood supply or passed on to the small intestine and bowel, with excess expelled as flatulence.

It has been reported that air residing in the pharynx when swallowed with food or saliva, is always swallowed after expiration of air from the lungs [32,33]. Inspired air from breathing contains ≈21% oxygen ≈78% nitrogen ≈1% argon and ≈0.004% carbon dioxide [34-36]. On expiration, air from the lungs contains ≈16% oxygen, ≈78% nitrogen, ≈1% argon and ≈5% carbon dioxide [34,35]. Therefore, swallowed air being expired air, must also contain 16% oxygen, 5% carbon dioxide and ≈1% argon. Note the water vapour contained in air has been excluded from the calculations.

The moles percent ratio of nitrogen : argon in ambient air has been reported as N_2_:Ar ≈ 83 : 1 (presumably calculated as N_2_:Ar ≈ 78/0.94) compared to that from flatulence of N_2_:Ar ≈ 72 : 1, slightly lower than that in air [37]. The nitrogen to oxygen ratio measured from flatulence was reported to be N_2_:O_2_ ≈ 11.5 :1.2, equivalent to N_2_:O_2_ ≈ 78 : 8.1 which indicates a ≈ 8 % lower value for oxygen compared to swallowed expired air of N_2_:O_2_ ≈ 78 :16 [37]. This indicates ≈ 8% of oxygen may have been used in the digestive system from the original 16% available from swallowed air.

### 2.10 Quantification of the volume of oxygen, carbon dioxide and nitrogen swallowed per 24 hours

From the calculated volume of air swallowed over 24 hours ≈ 6,435(315-46,688) ml, the volumes of the swallowed gases can be determined, with a large range of values shown, Table 7.

**Table 7.** The volumes of the major 3 gases that enter the digestive system through air swallowing /24 hours from the calculated volume of air swallowed of 6,435(315-46,688) ml and the recognition that swallowed air is from the expired air from the lungs with a high concentration of carbon dioxide, compared to the very low concentration in inspired air. Water vapour excluded from the calculations. Calculation methods in Appendix 1.

| Swallowed expired air volume 6435(315-46,688) ml | Average and range of swallow volume ml /24 hours |
| --- | --- |
| Volume of swallowed nitrogen ml /24 hours at 78% | 5000(250-36,400) |
| Volume of swallowed oxygen ml /24 hours at 16% | 1,000(50-7,500) |
| Volume of swallowed carbon dioxide ml /24 hours at 5% | 320(16-2,300) |
| Volume of swallowed argon ml /24 hours at 1% | 64(3-470) |

## 3. Discussion

### 3.1 The volume of air swallowed and digestive reflux disease

From the microphone method it was determined that while EDS there were 1 swallow ≈ 0.34 /minute (≈2.9 swallows /minute), while asleep ≈1 swallow /11.5 minutes (≈0.087 swallows /minute) and OTA ≈1 swallow every 2.6 minutes (≈ 0.39 swallows /minute), Table 1. The swallow rates /minute values correspond to air swallow rates when EDS ≈ 31 ml /minute, while asleep ≈ 1ml /minute and for OTA ≈ 4.3 ml /minute, Table 1. If complete aerobic digestion of food was related to the volume of air swallowed /minute and completed aerobic digestion reduced the probability of reflux diseases, then digestive reflux would be least likely to occur when most air is available, which occurs while EDS followed by OTA and finally most likely to occur when least air is available when asleep, as is often the case.

It has been reported the consumption of food less than 3 hours before sleep can be associated with an increased risk of GERD compared to patients with 4 or more hours before sleep [38]. Sleep alters respiratory breathing with rapid eye movement (REM) sleep having the highest number of breathing irregularity in both the frequency and respiratory rate where almost all body muscles, including respiratory muscles, become hypotonic except for the diaphragm, relied on to maintain tidal volume [33,39]. During sleep, irregular breathing and reduced air swallowing could create conditions unable to support luminal oxygen supply for aerobic digestion increasing the probability of developing reflux diseases [8].

Air swallowing has been related to digestive reflux diseases, with PPI refractory patients found to swallow more air ≈ 1.05±0.15 ml /minute while prandial than those who respond to PPIs ≈ 0.59±0.08 ml /minute [15]. It has been suggested that reduced air swallowing may be beneficial for some patient groups with GERD [15]. GERD patients who had used PPIs as a treatment, had stimulated (not prandial) saliva swallow volumes ≈ 1.3(0.7-2.0) ml /minute significantly higher than patients using polypharmacy (stimulated but not prandial) with volumes ≈ 0.967 ± 0.433 ml /minute, Table 5 [28,29]. Low saliva volumes, oral dryness and reduced postprandial swallowing frequency, have also been associated with GERD [40].

It would appear there is an optimal amount of air swallowing required to balance the volume of EDS and types of food consumed, to optimise gut electrochemistry and saliva swallowing, to provide numerous benefits including bicarbonate ions for acid buffering of the esophagus and stomach, with fluctuations likely to contribute towards developing digestive reflux [8,40,41].

The total volume of air swallowed calculated from the microphone method ≈ 6,400(31-47,000) ml /24 hours is quite small compared to the volume of air from breathing, which for an adult at rest is ≈ 8,640,000 ml /24 hours or ≈ 6,000 ml /minute air ventilation which with moderate exercise can increase to ≈ 40,000 ml /minute ventilation, Table 2 [35]. It is still not clear whether luminal oxygen supply from air swallowing, or luminal (and systemic) oxygen supply from breathing, is the major source of luminal oxygen for aerobic digestion. If air swallowing is the major source of luminal oxygen supply for aerobic digestion, then air swallowing is likely a vital factor for digestive health.

### 3.2 Limitations of the findings

It is assumed the volume of air swallows /bolus, determined from swallowing barium boluses, is equivalent to the volume of air swallowed per food or saliva bolus while EDS, asleep and at OTA. It is also assumed air is swallowed with each bolus (food, liquid or saliva) even though it is known some swallows are liquid only swallows, Table 3. Saliva boluses at OTA have a calculated swallow volume per swallow ≈ 0.56 ml with ≈ 1 swallow each minute or 0.56 ml /minute, Table 4,5 [30]. It has been reported that for 5ml barium swallows or even for saliva only swallows, the volume of air swallowed ≈14-15 ml [9,10]. It has also been shown that small bolus volumes can allow a greater air swallow volume than large volume boluses, Fig. 1. If less air is swallowed when barium or food boluses are absent and only saliva boluses are present when asleep, or at OTA, the lower volume of 0-1.7 ml /air swallows may reflect the true values.

Chewing food can allow oxygen from air to bind both chemically and physically to the food on forming the food bolus, increasing the air content of the stomach and adding to the volume of air swallowed but this additional air swallowed was not taken into consideration.

The range in the measured number of swallows ≈ 585(185-1459) /24 hours and the amount of air swallowed /bolus (11(1.7-32 ml)) have given a large range in the calculated air swallow volume ≈ 6,400(320-47,000) ml /24 hours which may not necessarily be an error of measurement but the possible variability that could be expected in the human population [16,35].

### 3.3 Future directions for GERD and digestive health studies

Aerophagia refers to excessive air swallowing and has been associated with symptoms of bloating and burping indicating air swallowing in general, can be undesirable. A more accurate description for the volume of air swallowed could be; excessive or mega aerophagia, healthy or adequate aerophagia or aerophagia and micro aerophagia for reduced air swallowing. A term for reduced air swallowing was not found presently in use and health conditions related to reduced air swallowing were also not found identified. Health conditions potentially related to micro aerophagia could result from reduced air swallowing at night, including nocturnal laryngopharyngeal and gastric reflux together with associated diseases like morning hoarseness of voice and morning halitosis found for GERD patients, possibly due to escaping odorous gas produced from inadequately aerobically digested food [8,43,44]. Further research into the roles of the swallowed gases oxygen, nitrogen and carbon dioxide in digestive health could be undertaken.

## 4. Conclusion

Oxygen supply to the gut lumen has a central role in aerobic digestion. Four models to quantify the volume of air swallowed /bolus were described, finding volumes between 0-40 ml were possible with the average value and range ≈ 11(1.7-32) ml /bolus. Values for the number of swallows in each state (EDS, asleep, OTA) were found from the literature and it was assumed the number of swallows directly related to the number of air swallows. From the number of swallows in each state, the total air swallow volume of ≈ 6,400(320-47,000) ml /24 hours was calculated. If improved aerobic digestion reduces the probability of reflux, then reflux would be least likely to occur while EDS with the highest air swallow volume /minute followed by OTA and finally most likely to occur when asleep, when the least air is swallowed, as is often the case. Low swallow rates when asleep can result in reduced luminal oxygen supply for digestion and may contribute to the increased risk of nocturnal digestive reflux diseases particularly for short dinner to sleep times of less than 3 hours.

## Abbreviations

CT: computer tomography
EDS: eating drinking snacking
GERD: gastroesophageal reflux disease
MRI: magnetic resonance imaging
OTA: other times awake
PPIs: proton pump inhibitors.

## 1 Appendix

**Table 1.**
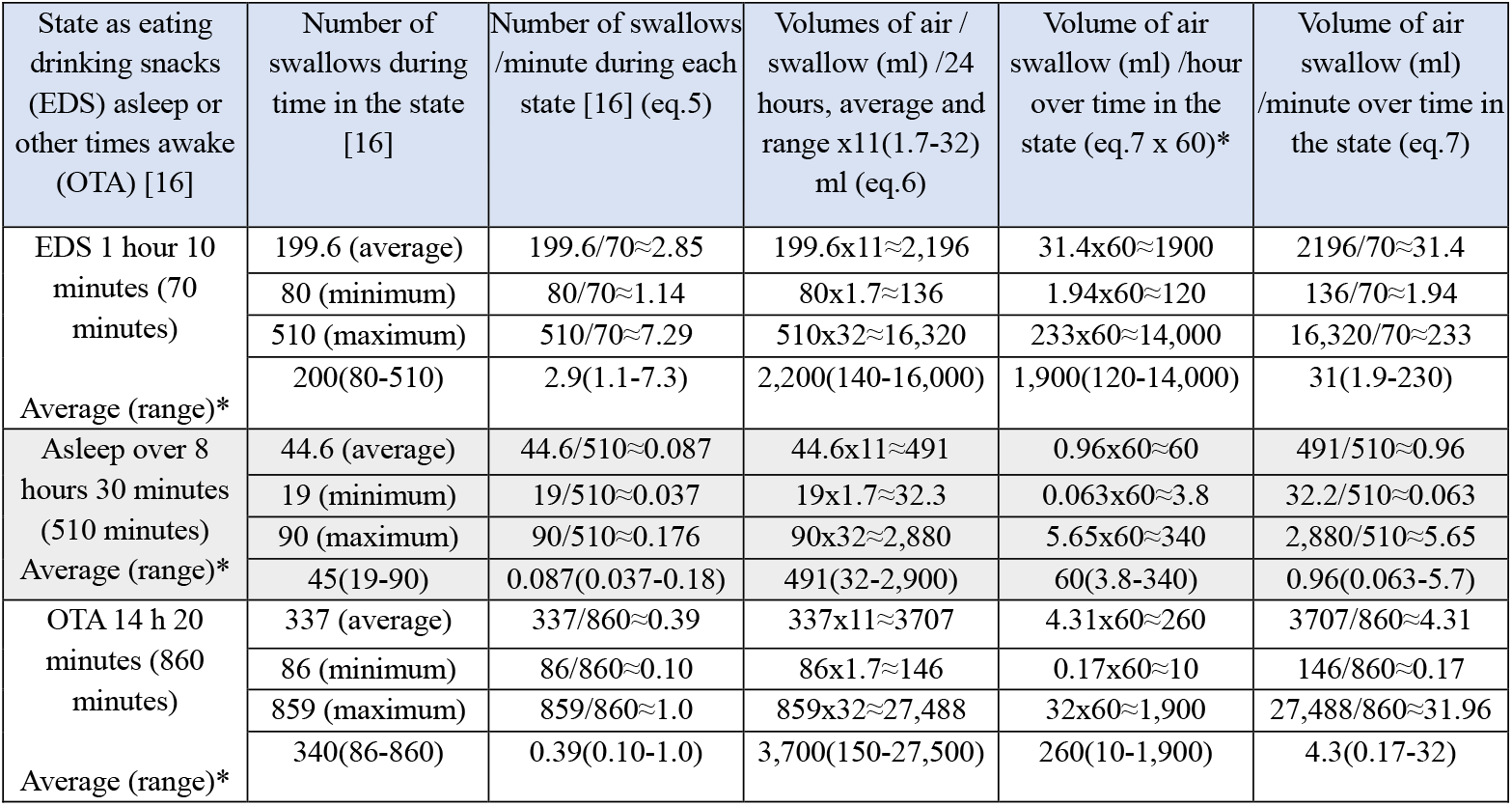
Calculation of air swallow volumes and rates from the number of swallows in each state [16].

**Table 2.**
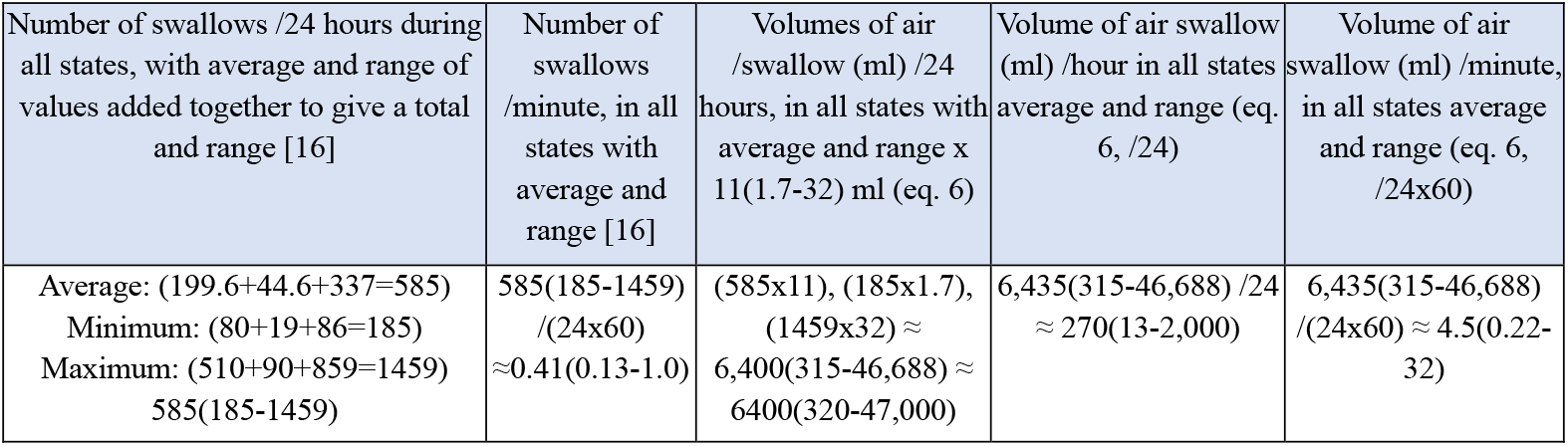
The number of swallows /minute and the volume of swallows /24 hours, /hours, and minutes in all states summed together [16].

**Table 7.** The volumes of the major 3 gases that enter the digestive system through air swallowing /24 hours from the calculated volume of air swallowed of 6,435(315-46,688) ml and the recognition that swallowed air is from the expired air from the lungs with a high concentration of carbon dioxide, compared to the very low concentration in inspired air. Nitrogen 78%, oxygen 16%, carbon dioxide 5%, argon 1%, with water vapour % excluded from the calculations.

| Swallowed expired air 6435(315-46,688) ml | Average swallow volume (ml) | Minimum swallow volume (ml) | Maximum swallow volume (ml) | Average and range of swallow volume (ml) |
| --- | --- | --- | --- | --- |
| Volume of nitrogen swallowed ml /24 hours | 6,435x0.78 ≈ 5020 | 315x0.78 ≈ 246 | 46,688x0.78 ≈ 36,400 | nitrogen 5000(250-36,400) ml /24 hours |
| Volume of oxygen swallowed ml /24 hours | 6,435x0.16 ≈ 1030 | 315x0.16 ≈ 50 | 46,688x0.16 ≈ 7500 | oxygen 1,000(50-7,500) ml /24 hours |
| Volume of carbon dioxide swallowed ml /24 hours | 6,435x0.05 ≈ 320 | 315x0.05 ≈ 16 | 46,688x0.05 ≈ 2,300 | carbon dioxide 320(16-2,300) ml /24 hours |
| Volume of argon swallowed ml /24 hours | 6435x0.01 ≈ 64 | 315x0.01 ≈ 3 | 46,688x0.01 ≈ 470 | argon 64(3-470) ml /24 hours |

